# Mitochondrial sequences or Numts – By-catch differs between sequencing methods

**DOI:** 10.1101/739805

**Authors:** Hannes Becher, Richard A Nichols

**Affiliations:** Queen Mary University of London, School of Biological and Chemical Sciences; University of Edinburgh, Institute of Evolutionary Biology

**Keywords:** mitochondrial genome, plastid, Numt content, Illumina, PacBio, tangle plot

## Abstract

Nuclear inserts derived from mitochondrial DNA (Numts) encode valuable information. Being mostly non-functional, and accumulating mutations more slowly than mitochondrial sequence, they act like molecular fossils – they preserve information on the ancestral sequences of the mitochondrial DNA. In addition, changes to the Numt sequence since their insertion into the nuclear genome carry information about the nuclear phylogeny. These attributes cannot be reliably exploited if Numt sequence is confused with the mitochondrial genome (mtDNA). The analysis of mtDNA would be similarly compromised by any confusion, for example producing misleading results in DNA barcoding that used mtDNA sequence. We propose a method to distinguish Numts from mtDNA, without the need for comprehensive assembly of the nuclear genome or the physical separation of organelles and nuclei. It exploits the different biases of long and short-read sequencing. We find that short-read data yield mainly mtDNA sequences, whereas long-read sequencing strongly enriches for Numt sequences. We demonstrate the method using genome-skimming (coverage < 1x) data obtained on Illumina short-read and PacBio long-read technology from DNA extracted from six grasshopper individuals. The mitochondrial genome sequences were assembled from the short-read data despite the presence of Numts. The PacBio data contained a much higher proportion of Numt reads (over 16-fold), making us caution against the use of long-read methods for studies using mitochondrial loci. We obtained two estimates of the genomic proportion of Numts. Finally, we introduce “tangle plots”, a way of visualising Numt structural rearrangements and comparing them between samples.

## Introduction

Sequences of mitochondrial DNA have proved indispensable markers for population genetics and phylogenetics for decades (Avise et al., 1987; Ballard & Rand, 2005). More recently, numerous ecological experiments have exploited the universal animal barcoding marker, COX1, which is a mitochondrial gene (Hebert, Ratnasingham, & deWaard, 2003). One valuable property of mitochondrial sequences is that, being more abundant, they tend to be more effectively amplified by PCR than nuclear sequences, in particular in difficult or degraded samples. Mitochondrial markers are therefore widely used in research on museum specimens (Anmarkrud & Lifjeld, 2017), ancient DNA studies (Baca et al., 2018; Mohandesan et al., 2017) and analyses of faecal samples (van der Valk, Lona Durazo, Dalén, & Guschanski, 2017). A second advantage is that, thanks to their comparatively small size (approximately 16 kbp) and conserved structure (Boore, 1999) animal mitochondrial genomes are easy to assemble. Similar considerations make plastid genomes particularly valuable in genetic analysis of plants (Twyford & Ness, 2016).

The advantages of mtDNA analysis can be negated by the presence of Numts: nuclear inserts derived from mitochondrial DNA (and in plants, the plastid equivalent, Nupts). Evidence of such insertions was first found shortly after mitochondria were discovered to contain their own genetic material (Du Buy & Riley, 1967) and it has since become clear that Numts are present in many species (Bensasson, Zhang, Hartl, & Hewitt, 2001), often in multiple copies. The abundance of Numts strongly depends on whether a species has one or more mitochondria per cell, an observation which led Barbrook, Howe, & Purton (2006) to postulate a limited transfer window.

The confusion of Numts and mitochondrial sequence could lead to incorrect interpretations of molecular genetics studies (Blacket, Semeraro, & Malipatil, 2012; Hawlitschek et al., 2017; Jordal & Kambestad, 2014; Kim, Lee, & Ju, 2013; Thalmann, Hebler, Poinar, Pääbo, & Vigilant, 2004; Zhang & Hewitt, 1997). Any study targeting mitochondrial sequences will therefore benefit from knowledge of the genomic content of Numts.

While Numts are commonly seen as a nuisance, they are fascinating study objects in their own right. Accruing substitutions more slowly than the mitochondrial DNA lineage, they act as “molecular fossils” providing information about the ancestral mitochondrial sequence (Lopez, Yuhki, Masuda, Modi, & O’Brien, 1994; Thalmann, Serre, et al., 2004). Some can be distinguished, because substitutions which have accumulated since integration into the nucleus have a high incidence of non-synonymous changes relative to mtDNA (Bensasson, Zhang, & Hewitt, 2000). Such Numts can be used as genetic markers, for example providing evidence of past episodes of hybridisation between taxa (Brelsford, Mila, & Irwin, 2011; Miraldo, Hewitt, Dear, Paulo, & Emerson, 2012).

Before high-throughput sequencing data became readily available, Numts could be detected, albeit with some difficulty, by PCR-based methods (Bensasson et al., 2000) or cytologically, by *in situ* hybridisation of mitochondrial sequences to chromosomal preparations (Gellissen, Bradfield, White, & Wyatt, 1983). To physically separate mitochondrial and nuclear DNA, (ultra) centrifugation can be used (Garber & Yoder, 1983; Lansman, Shade, Shapira, & Avise, 1981), but these methods require considerable technical effort. Since the advent of high-throughput sequencing data, two further approaches have been applied to identifying Numts. (1) In well-assembled genomes sequenced at high coverage, Numts can be detected simply by screening the assembly for regions with similarity to mitochondrial DNA (Hazkani-Covo, Zeller, & Martin, 2010). Where genomic data has been assembled for multiple individuals, as in humans, insertion/deletion polymorphism for Numts can be readily studied (Dayama, Emery, Kidd, & Mills, 2014). (2) In absence of a well-assembled genome, for instance in genome skimming studies (Dodsworth, 2015; Straub et al., 2012), Numts may be identified if reads (or read pairs) match the mitochondrial sequence for only part of their length. Otherwise, if the whole of the read maps to the mitochondrial genome, it may be possible to classify its Numt origin if the sequence has diverged from the mitochondria. Those that have not diverged sufficiently to be distinguished are termed “cryptic Numts” (Bertheau, Schuler, Krumböck, Arthofer, & Stauffer, 2011).

In this paper, we develop a further approach that investigates Numts by making use of the “by-catch” from High-throughput sequencing. We use this term to emphasise that much sequencing data is superfluous to the aims of a specific experiment, often the huge majority. The low price of sequencing data means these data could be discarded; yet this genomic by-catch can be mined for valuable incidental information. In particular it can be used to assemble genomes of organelles such as mitochondria. The choice of sequencing platform may influence the type of by-catch produced, particularly because of differences in fragmentation and size-selection protocols. We show that this difference can be exploited to investigate the Numt content of the genome.

In order to outline our approach, it is helpful to divide the data generated from a sequencing library into fractions (see Figure 1). The first major division is between A – reads without similarity to mitochondrial sequences and ML – mitochondrial-like sequences (i.e. those which align to the mitochondrial genome). This ML fraction may conceptually be subdivided further into D – those which can be identified as Numt sequences (i.e. sequences aligning for their full length but having a different sequence, or aligning for part of their length), C – cryptic Numts indistinguishable from actual mitochondrial sequences, and M – sequences derived from actual mitochondrial genomes. Fractions A, C, and D are derived from nuclear DNA (fraction N, comprising A + C + D). In our analysis we assume that their ratios are effectively constant among individuals from the same species. For instance, the proportion of obvious Numts in the nuclear genome D/N should be constant. The size of fraction M, which is contributed by mitochondria, may differ between samples because of differences in the number of mitochondria per cell with tissue, sex, or developmental stage (Fernández-Vizarra, Enríquez, Pérez-Martos, Montoya, & Fernández-Silva, 2011). This will cause difference in M among samples, which will be observed in differences in the ratios of A and ML in different individual samples.

**Figure 1.**
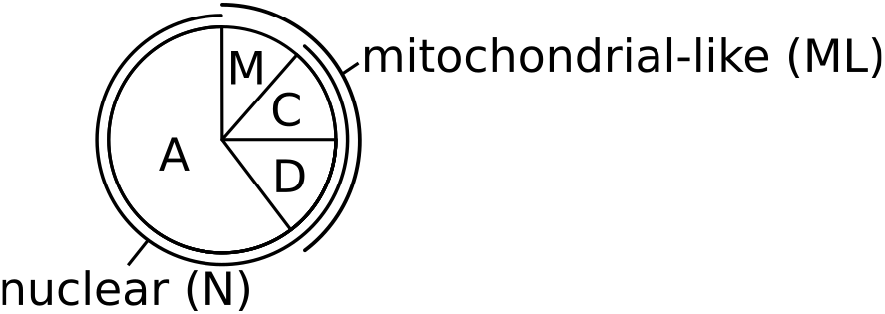
Fractionation of sequencing data. Of all the sequencing data generated from one sample, some fraction N will originate from the nuclear DNA and some fraction M from the mitochondrial DNA, and other sequences. The nuclear fraction N may contain Numts. Some of these will have sequences divergent from M and hence are easily identifiable (fraction D). Other Numts may be cryptic, e.g. recent insertions, (fraction C). Together, Numts and mitochondrial sequences for the fraction ML “mitochondrial-like”.

Here we demonstrate two complementary approaches for estimating the proportion the nuclear genome made up of Numts. One exploits the variation in M from sample to sample in short-read data, which arises because of differences in the mitochondrial composition with tissue, sex, or developmental stage. Secondly in some types of long-read data M is minimal, so the ratio can be calculated directly. We demonstrate these approaches using genome skimming data (coverage < 1/3x) generated by short-read (Illumina’s NextSeq) and long-read (PacBio’s RSII) platforms from the grasshopper, *Podisma pedestris*. We assemble the species’s mitochondrial genome sequence and calculate these two estimates of the proportion of Numts in the nuclear genome. We also introduce a method, which we call “tangle plots”, for the visualisation of Numts with structural re-arrangements

## Materials and Methods

### Samples and sequencing

Information about the samples can be found in Table 1.

**Table 1.** Overview of the sequencing data generated and fraction ML (mitochondrial-like, mapping to the mitochondrial genome assembly).

| NextSeq | Sample | Origin | High-quality<br>reads | ML (% of<br>reads) | Max SNP allele<br>frequency |
| --- | --- | --- | --- | --- | --- |
|  | N1 | Blayeul | 25600134 | 0.48% | 15% |
|  | N2 | Blayeul | 37563526 | 0.64% | 14% |
|  | N3 | Blayeul | 13034544 | 1.17% | 8% |
|  | N4 | Mariaud | 35257122 | 0.97% | 9% |
|  | N5 | Mariaud | 33393734 | 1.32% | 7% |
|  | N6 | Mariaud | 21813470 | 0.44% | 20% |

| PacBio | Sample | Origin | Reads (bp in CCS) | ML reads (bp) |
| --- | --- | --- | --- | --- |
|  | PB1 | Blayeul | 92971 (336152778 bp) | 160 (0.04%) |
|  | PB2 | Blayeul | 97937 (376785844 bp) | 131 (0.03%) |
|  | PB3 | Bournee | 106991 (248262810 bp) | 152 (0.06%) |

### Illumina NextSeq

Freshly removed hindlegs of *Podisma pedestris* were snap-frozen and stored at −79 °C. Before DNA extraction, the legs were dipped into boiling water to inactivate DNases. Subsequently, the denatured femur muscle was dissected out. DNA was then extracted using a Qiagen Blood and Tissue kit following the manufacturer’s instructions. Using a Covaris ultra sonicator the DNA was sheared aiming to achieve a median size of 550 bp. Libraries for sequencing were prepared using an Illumina TruSeq DNA PCR-Free kit. Sequencing was carried out at QMUL’s Genome Centre on Illumina’s NextSeq Platform using v2 chemistry.

### PacBio RSII

Freshly removed hindlegs were stored in pure ethanol. DNA was extracted from four samples using a Qiagen Gentra HMW kit resulting in molecules with a length of mainly > 48 kbp (TapeStation, Agilent Genomics). Further work was carried out by The University of Liverpool’s Centre for Genomic Research. The aimed size for DNA fragmentation was 10 kbp. The libraries’ median (non-redundant) insert sizes were 3125, 3167, and 2097 bp. Sequencing was carried out on a PacBio RSII machine using C6 chemistry. All PacBio data were obtained as circular consensus sequences (CSSs) in FASTQ format. These are of a higher per-base quality than the raw reads, because they are generated from multiple reads generated from the same circular template.

### Data cleaning

Two sets of clean NextSeq data were prepared. For the RepeatExplorer analysis (see below), the data were filtered using a custom python script keeping only read pairs where 90 % of the bases had a phred quality score > 20. Pairs matching the TruSeq adapters (detected by BLASTn num_aligments 1) were discarded to remove adapter dimers.

A second cleaned set of NextSeq data was generated for mapping and variant calling. Here, we aimed to remove as many low-quality base calls as possible. The first 5 bp of each read were removed and, using Skewer (Jiang, Lei, Ding, & Zhu, 2014), each 3’ end was trimmed until the last base had a phred quality 30 or higher.

For the RepeatExplorer analysis of PacBio data, pseudo paired reads of 151 bp with an insert size of 550 bp were cut out of long PacBio reads using custom-made python scripts which depend on the biopython module, http://biopython.org/ (Cock et al., 2009).

### RepeatExplorer analyses

RepeatExplorer (https://galaxy-elixir.cerit-sc.cz/, Novák, Neumann, Pech, Steinhaisl, & Macas, 2013) is a pipeline for analysing the repetitive genome content from short-read genome skimming data. It performs an all-to-all comparison and generates clusters of similar reads, which often correspond to particular genomic repeats such as transposable elements or satellites. Mitochondrial genomes and ribosomal RNA genes (rDNA), which are present in high copy numbers, are usually picked up as well.

RepeatExplorer was run twice. The first run was used to assemble a reference sequence for the *Podisma pedestris* mitochondrial genome, from the NextSeq reads. In the second run, 100,000 NextSeq read pairs from each of six individuals (N1-N6) and 150,000 PacBio pseudo read pairs from each of three individuals (P1-P3) were analysed jointly to compare the sequencing methods. The pipeline was supplied with a custom annotation database containing the mitochondrial genome sequence of *Schistocerca gregaria* [Genebank NC_013240.1 (Erler, Ferenz, Moritz, & Kaatz, 2010)] in the first round and with the *Podisma pedestris* rDNA and mitochondrial genome in the second run.

### Mitochondrial sequence assembly

Eight RepeatExplorer clusters connected by paired reads (244 - 57 - 230 - 205 - 69 - 85 - 102 - 161) showed sequence similarity to *S. gregaria* mitochondrial DNA. Those clusters overlapping consensus sequences were assembled in Geneious R9, forming a reference to which reads of sample N1 were mapped. High coverage and truncated reads at the control region indicated a duplication, which was then added to the reference. Subsequently, each of the six NextSeq samples’ sequencing data were mapped individually using BWA (Li & Durbin, 2009) with the following command line: `bwa mem -t <NO of threads> <REFERENCE> <(zcat read files)`. For each of the six alignments, 50 % majority rule consensus sequences were created in Geneious. They were annotated automatically using the MITOS WebServer (Version 2 beta, Bernt et al., 2013). Our mitochondrial genome assemblies were checked by re-assembling the short-read data with NOVOPlasty (Dierckxsens, Mardulyn, & Smits, 2016), which produced essentially the same sequences.

## Mapping and variant detection

In order to detect individual-specific variants, a second round of mapping was carried out with NextSeq data. Polymorphisms were called using Geneious’s function “Find Variations/SNPs” with default settings and a minimum allele frequency set to 0.01. The resulting tables were exported to CSV format and were processed interactively in R 3.3.1 (R Core Team, 2016).

All PacBio reads were aligned to the mitochondrial assembly using the LAST suite (Kielbasa, Wan, Sato, Horton, & Frith, 2011). In brief, the reference genome was masked in regions with GC-content below 10 % and was subsequently converted to a LAST database using the scoring scheme NEAR, preserving all masked regions and additionally masking simple repeats (optimised for high AT-content). Lastal was then run with parameter D set to one thousand times the length of the assembly (corresponding to an e-value of 1e-3 in BLAST). Of the resulting hits, only those with alignment lengths above 100 bp were kept. Shorter ones tended to map to in regions of low complexity, not permitting meaningful conclusions about homology.

### Assessment of sequencing bias and genomic proportion of Numts

The output of the comparative (i.e. second) RepeatExplorer run was used to assess sequencing technology-specific bias (see Fig. 2).

**Figure 2.**
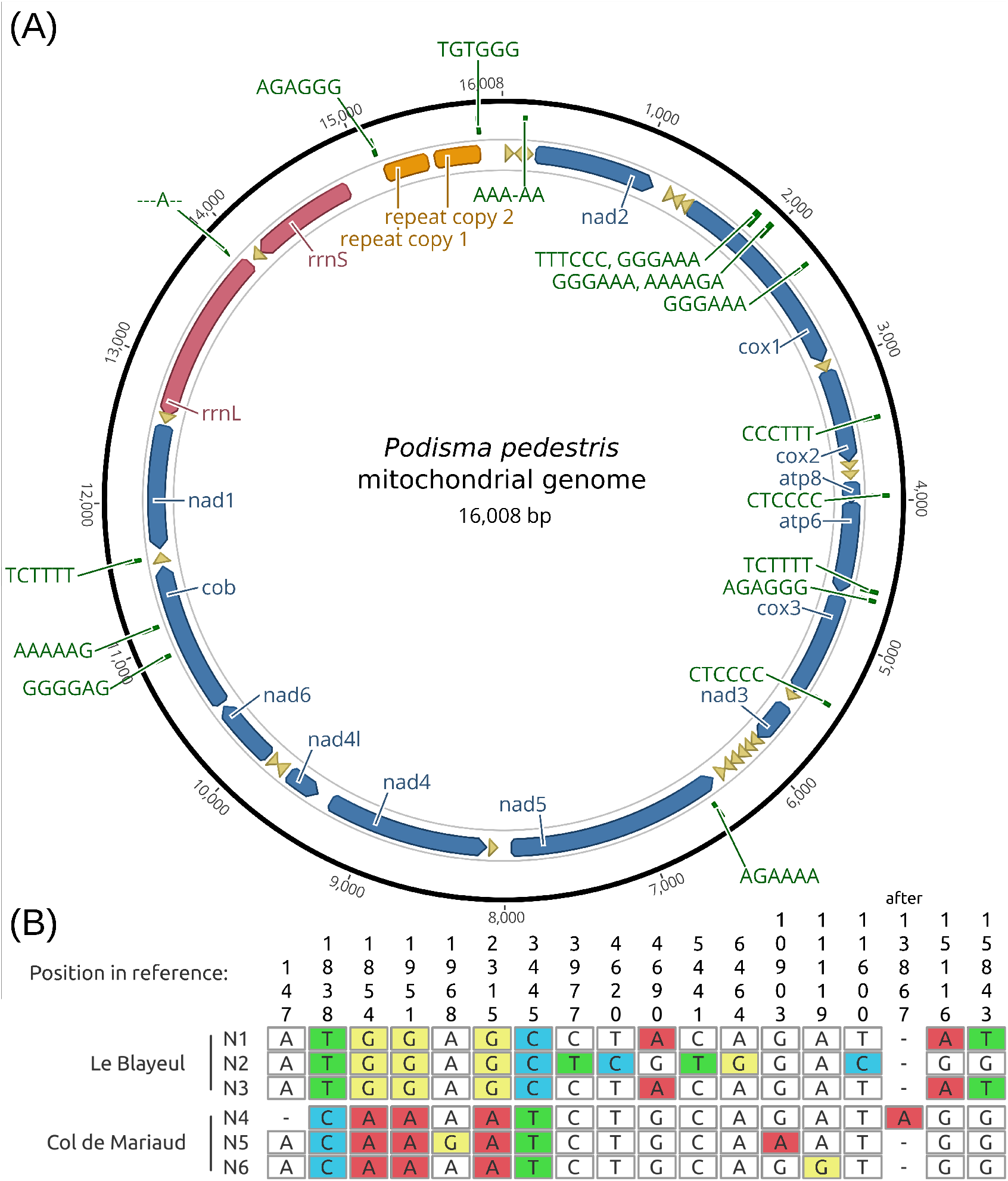
Annotated mitochondrial genome assembly of the alpine grasshopper, *Podisma pedestris*, and between-individual polymorphisms. (A) Shows a representation of the mitochondrial genome assembly. (B) The alignment of the six assemblies contains 18 polymorphic sites. Their positions are indicated above the alignment and in (A). Mismatches are highlighted.

## Illumina data

Sequencing reads of the mitochondrial-like fraction (ML, see Fig. 1) could either have originated from mitochondrial genomes (M) or from Numts (fractions C and D). Assuming each of the individuals contained the same genomic proportion of Numts, the variation between samples in the relative proportion of ML in the NextSeq data (see Tab. 1) would be attributed to different mitochondrial densities in the extracts (varying proportion of M). One estimate of the proportion of Numt sequences reads can be obtained from the assumption that most mtDNA in any one individual is monomorphic, whereas some of the Numt sequences will be fixed for a different allele (because mtDNA tends to evolve faster than Numts). In this case, the frequency of this Numt allele will be proportional to the relative contribution of Numts. The maximum of the distribution of allele frequencies (shown as horizontal bands of dots in Fig. 4) provides an estimate for the relative contribution of Numts to the data mapping to each mitochondrial assembly.

This assumption is supported by the very close correlation (R^2^=0.93, p=0.001) between the maximum allele frequency and the proportion of reads (A) which are not mitochondrial-like.

### PacBio data

The estimate of M from the PacBio data was estimated as follows. CCSs aligning only partially (< 95%) were considered Numt-derived. Reads matching along > 95 % of their length were considered full-length matches. For these, alignment error profiles were compared to the reads’ phred quality scores. If an alignment contained significantly more mismatches than expected (5 % confidence interval, one-sided, Bonferroni-corrected for 59 alignments), it was considered a Numt sequence. The remainder of the full-length matches were provisionally classified as mitochondrial, belonging to fraction M.

### Tangle plots

Code to reproduce the example shown in figure 5, as well as explanations, and distance computation can be found in the GitHub repository “tangles”. See also supplemental information (https://github.com/SBCSnicholsLab/tangles), which contain explanations and another example.

## Results

### Six mitochondrial genome assemblies

We assembled the mitochondrial genome sequences of six individuals of *Podisma pedestris* (each of our short-read genome-skimming datasets) using contigs produced by the RepeatExplorer pipeline (Novák et al., 2013). RepeatExplorer generates “clusters” of decreasing size corresponding to repetitive DNA sequences in the samples analysed. RepeatExplorer contigs with similarity to the mitochondrial genome of the desert locust, *Schistocerca gregaria*, were merged in Geneious R9 and a 383-pb direct repeat, which had been collapsed, was adjusted manually after mapping each sample’s reads back to the respective assembly. To check the reliability of this approach, we re-assembled the mitochondrial genome from each data set with NOVOPlasty, yielding essentially the same sequences, the differences being around the control region; for example NOVOPlasty did not assemble the repeat in 3 cases.

Each of our assemblies is 16,008 bp in length. The average mapping depth varies between samples from several hundred to few thousand-fold, which could be due to differences in cellular content of mitochondria between individuals (Tab. 1). All genes typically found in animals were identified using the MITOS WebServer v2beta (Fig. 3A). The gene order is collinear with other grasshopper mitochondrial genomes, and the sequences align readily (see alignment in supplementary file S1).

**Figure 3.**
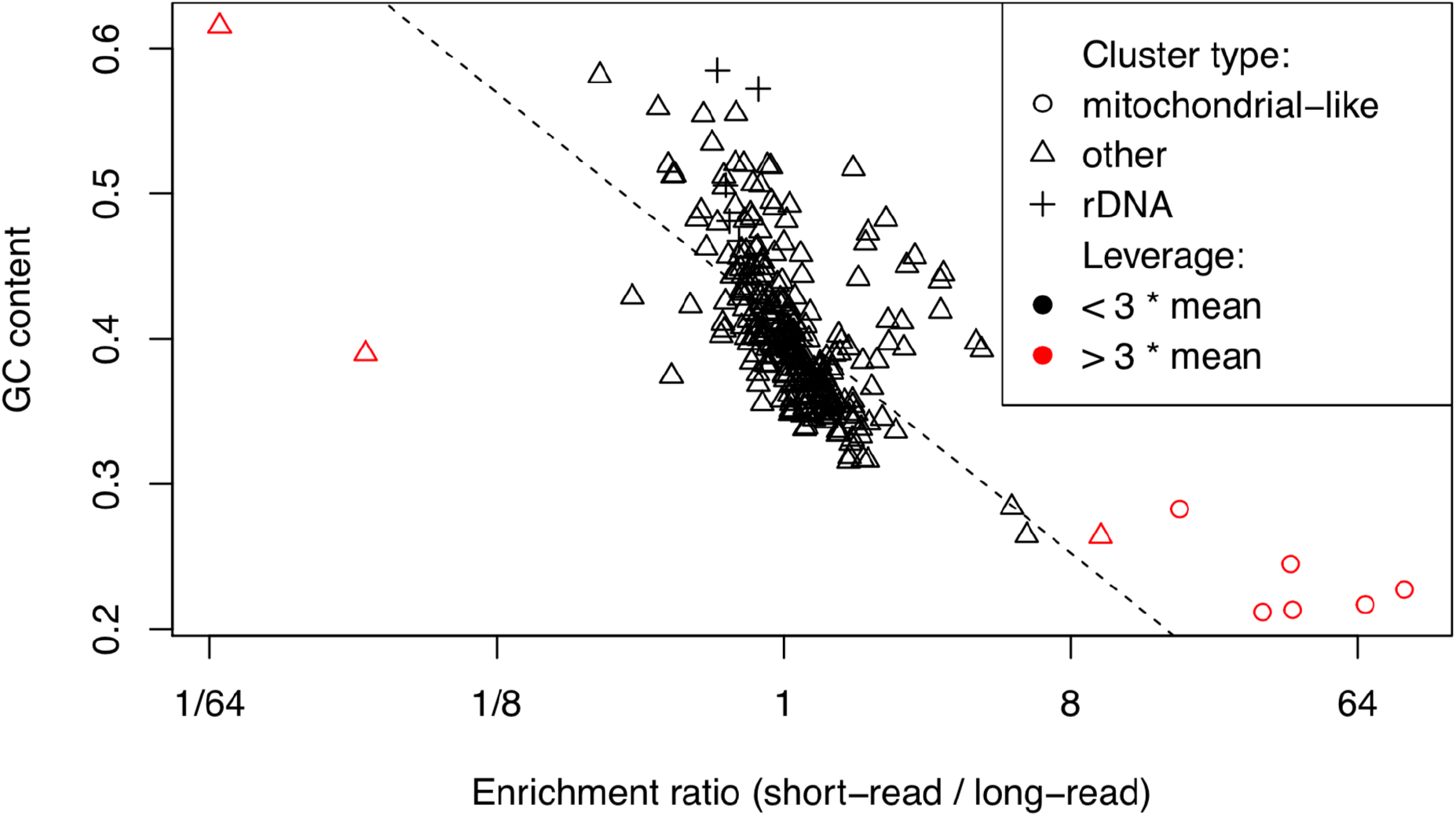
Enrichment for short or long reads plotted against GC-content for 300 RepeatExplorer clusters. Symbols right of the centre indicate high-abundance DNAs enriched in the short-read data. There is a general trend for enrichment in PacBio data for clusters with higher GC-content (regression line). Mitochondrial-like clusters, which show the strongest bias, are indicated as circles, rDNA as crosses. Data points with a leverage greater than three times the mean leverage are shown in red. These were excluded from the final regression (dashed line).

The alignment of all six consensus sequences contains 18 variable sites, five of which show population-specific polymorphisms (Fig. 3B). A neighbour-joining tree shows that each population’s individuals have sequences most similar to one other (see tree in supplementary data S2). The PacBio data did not yield enough mitochondria-like sequence to attempt an assembly, see below.

### The amount of ML data differs between sequencing methods

The proportion of each dataset with similarity to mitochondrial sequences (mitochondrial-like, ML in Fig. 1) was identified by mapping reads back to the assembly (of sample N1). It can be seen from Tab. 1 that the ML fraction in short read data (proportion of reads) is at least one order of magnitude larger than the ML fraction in long-read data (sum of read lengths) in all samples.

### High-abundance sequences differ between Illumina and PacBio data

Because of the comparatively low coverage of the genome skimming data generated, it is only possible to compare sequences that are very abundant in the libraries sequenced, such as genomic repeats and organelle DNA. In order to compare the data generated by PacBio and Illumina sequencing, we used RepeatExplorer; an pipeline that generates clusters corresponding to high-abundance sequences. If both Illumina and PacBio sequencing were unbiased in their representation of the DNA found in our samples, then each cluster should contain similar proportions of PacBio and Illumina data, corresponding to the amount of data put into RE. The proportion of short-read data in each of the 300 largest clusters is shown in Fig. 3 (on a logarithmic scale). As is commonly seen, the short-read data contain fewer sequences with higher GC content (Ekblom, Smeds, & Ellegren, 2014). While this bias is between ¼ and 4-fold for most clusters, the mitochondrial-like clusters (circles in Fig. 3) show the most extreme values (at least 16-fold enrichment in our short-read data).

### Short reads: Polymorphism in individual-specific alignments

SNPs were called within each alignment of individual-specific short ML reads. The minimum minor allele frequency set to 1 % to avoid erroneous calls due to sequencing errors. All assemblies contain numerous polymorphic sites with low to medium minor allele frequencies, which can be interpreted as variants present in Numts (dots in Fig. 4). The fact that we find appreciable allele frequencies even though we sequenced only a fraction of each genome, strongly suggests that there a multiple Numt insertions present in each sample. The distributions of these allele frequencies are skewed towards 0 with maxima varying between samples (the extremes are 7 % and 20 % in samples N5 and N6, corresponding to the narrowest and widest band in Fig. 4). Over all samples, there is a correlation between fraction D (distinguishable Numts) and fraction A (without sequence similarity to mitochondrial genomes). The slope of the linear regression represents the genomic proportion of distinguishable Numts in *P. pedestris*. It is 9×10^-04^ (p=1.04×10^-3^) with a standard error of 1×10^-4^. As expected, the intercept is not significantly different from zero (p=0.805), see Supplemental figure S6.

**Figure 4.**
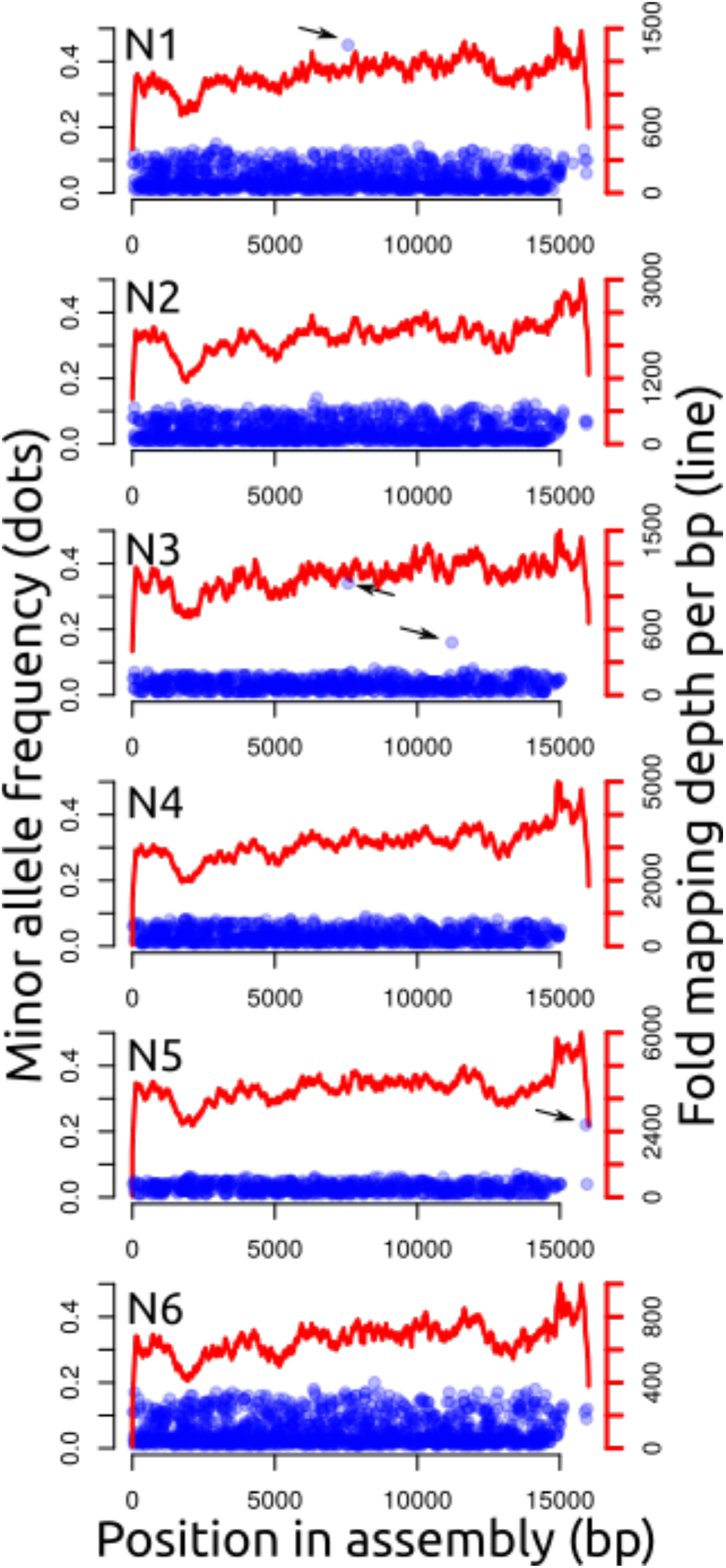
Polymorphisms within alignments. For each individual, the minor allele frequencies and positions of SNPs are shown as dots. Note, the dots generally form bands with widths differing between samples. There are few outliers, marked with arrows, likely indicating heteroplasmy. Individuals N1 and N3 share such a polymorphism at base pair 7567. The lines indicate per-base pair mapping depths.

In total, there are four SNPs with frequencies that are clear outliers from the frequency distribution (shown by arrows in Fig. 4). Interestingly, individuals N1 and N3 from Le Blayeul share one such polymorphism at base pair 7567. These high-frequency variants are presumably the signatures of heteroplasmy (as seen by Mao et al., 2014 in bats).

### Tangle plots: Atypical distances between read-pairs as a signature of Numts

Paired reads mapped to the mitochondrial assembly could have originated from mitochondrial genomes (M fraction) or from Numts. Those from the M fraction should show intra-pair distances consistent with the libraries’ insert sizes, since the mitochondrial genome is highly conserved. Conversely, Numt sequences may have been subject to insertions, deletions, or rearrangements resulting in longer distances between map locations on the mitochondrial genome or discordant read orientations. The majority of intra-pair differences fell into a distribution with a maximum around 400 bp representing the sequencing insert size (supplemental data S3). There is a second (much shallower) peak above 15,000 bp resulting from mapping reads generated from circular molecule to a linearised reference.

There is a small subset of read pairs with intermediate mapping distances that might be attributed to Numt sequences containing deletions or rearrangements. Fig. 5 shows each of these intermediate read pairs (those with an intra-pair distance between 1500 bp 14,508 bp) as a line connecting the paired reads’ positions resulting in “tangle plots”. Interestingly, some of the lines shown cluster together. Given the low sequencing coverage, this strongly suggests the presence of multiple copies of some Numts. Some patterns are shared across multiple samples, but the overall patterns are not population-specific (a linear discriminant analysis failed to assign all individuals to the correct populations, not shown here).

**Figure 5.**
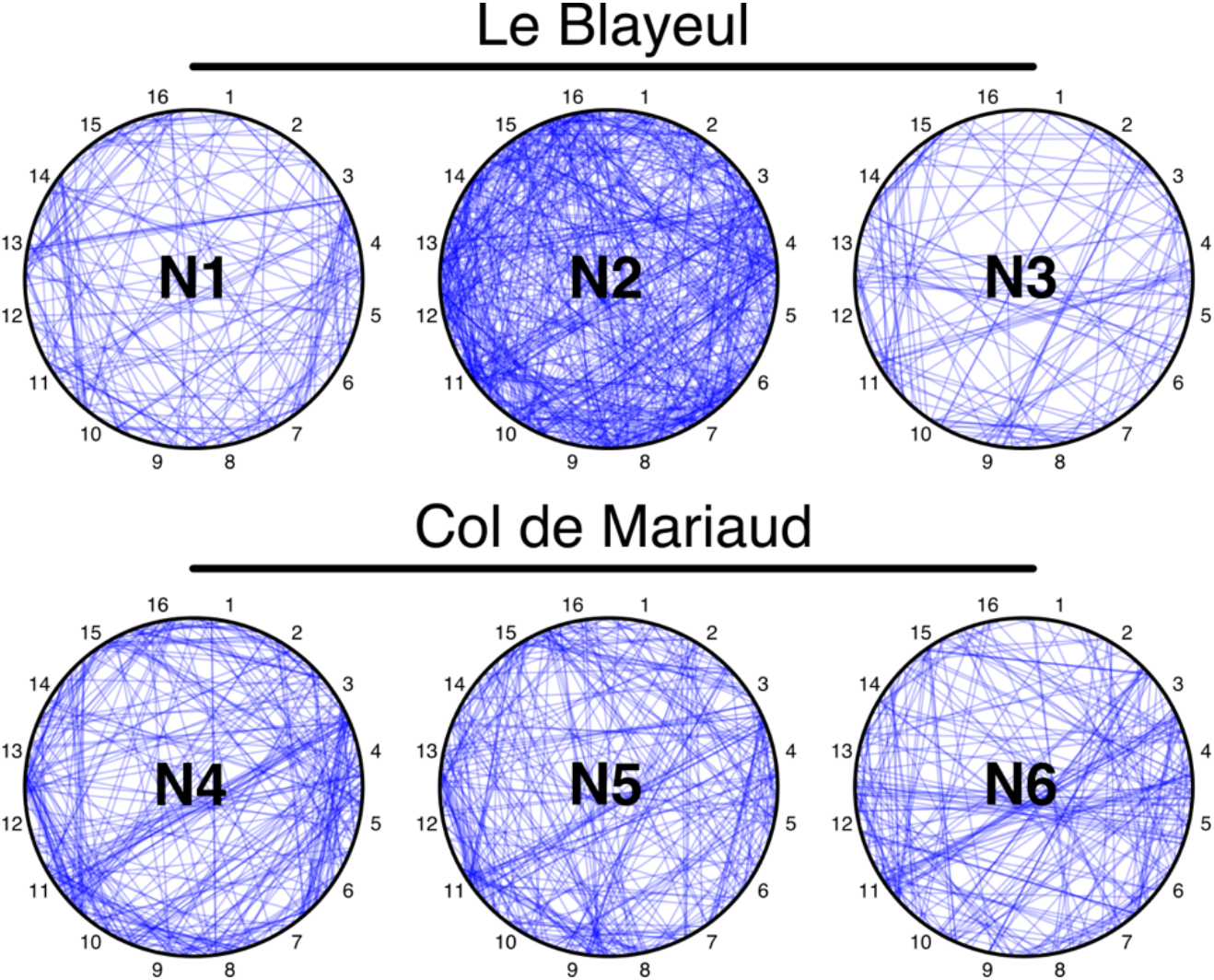
Tangle plots. The position of read pairs mapped with unusual read distances is indicated by a line connecting the read locations. For ease of comparison, 16 positions are labelled on each graph. Note common patterns between individuals. For instance, all individuals in the Col de Mariaud population show several read pairs connecting positions 3/4 and 11. All samples of both populations show read pairs connecting segments 3/4 and 7.

### Mapping PacBio reads

PacBio circular consensus sequences (CCS) generated from DNA of three individuals (samples P1-P3) were mapped to the mitochondrial assembly. Out of 297,899 non-redundant reads generated in total, 443 showed similarity to the mitochondrial assembly with a cumulative mapping length of 396,770 bp. Of these, most reads matched the mitochondrial reference only along a part of their length, a pattern expected for short Numts and also chimeric PacBio read (which we expect to be rare). The alignments cover 263,635 bp, which corresponds to 0.027 % of the total CCS data generated. There were only 59 PacBio CCSs matching full-length, of which 41 were sufficiently diverged from the mitochondrial sequence to meet or criterion for classification as Numts. These align along 96,210 bp corresponding to 0.01 % of the (non-redundant) PacBio data generated. The remaining 18 full-length matches could be derived from mitochondrial genomes, but they may well be derived from Numts inserted recently. Covering 36,925 bp, these ambiguous CCSs represent only 9.3 % of the ML fraction in the PacBio data.

Interestingly, the mapping depth of full-length matches has a bimodal distribution. While the 18 ambiguous matches contribute mostly to the first peak, Numt-derived CCSs map to the areas under both peaks (see Fig. 6).

**Figure 6.**
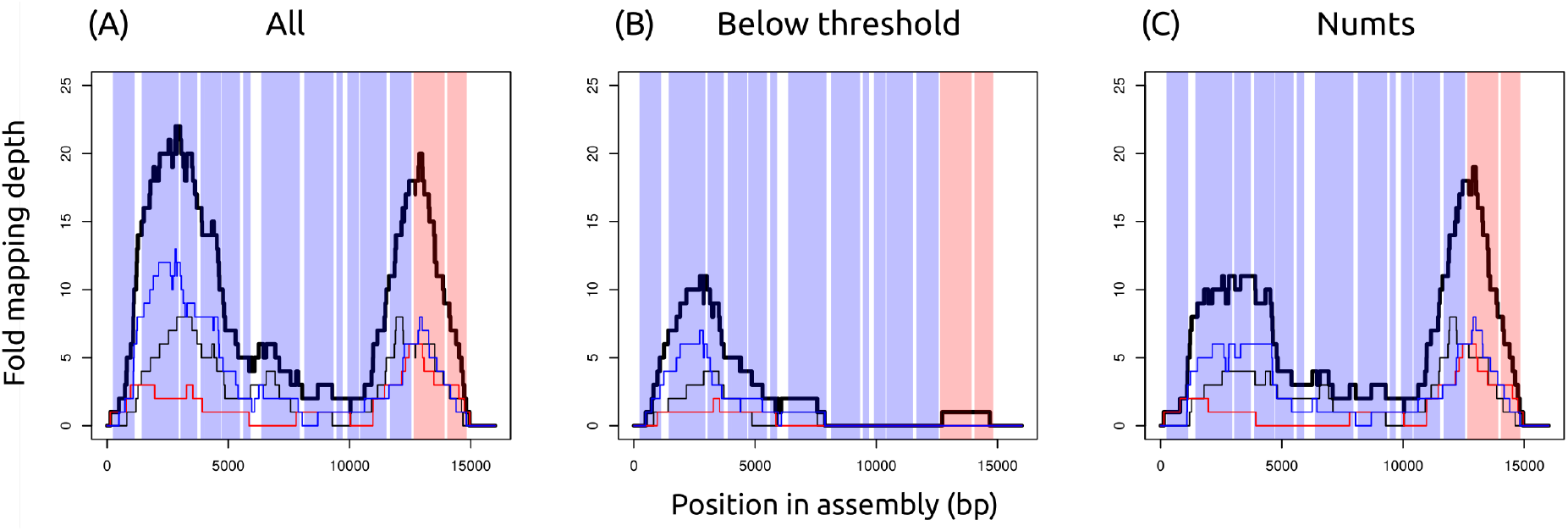
Mapping depths of PacBio CCSs mapping to mitochondria for their full-length. The bold line indicates the total over all three samples, narrow lines represent individual samples. Vertical shaded bands indicate the positions of protein-coding genes and mitochondrial rRNA genes (the two salmon coloured bands on the far right). (A) All CCSs matching full-length. (B) Fraction M or C (either mitochondrial or cryptic Numt) (C) Fraction D (classified as a Numt).

## Discussion

Both mitochondrial (M) and Numt (C+D) sequence are generated as side-products of sequencing experiments, analgous to by-catch on fishing trawlers. We investigated these sequences using genome-skimming data (less than ⅓x genomic coverage) from the grasshopper, *Podisma pedestris*, using Illumina’s NextSeq and PacBio’s RSII platforms with six and three biological replicates, respectively

### By-catch differs between sequencing methods

One of the most striking results is that the Illumina data had over 16-fold higher frequency of reads mapping to the mitochondrial clusters than the PacBio data, suggesting that the Illumina protocol produced a correspondingly higher proportion of sequences from the mitochondria (the M fraction), at some point between extraction and data interpretation. This bias cannot be explained by the known general over-representation of sequences with low-GC sequences in Illumina reads (Ekblom et al., 2014), as shown by the deviation of the mitochondrial clusters from the general trend in Fig. 3. The result is reinforced by the comparable bias shown in the estimates of the proportion of mitochondrial sequences classified as Numts (fraction D/ML). This value is also much higher in the PacBio data than the Illumina. In the PacBio case, the two D categories sum to 91% of ML (the D estimate is obtained from the length of the mitochondrial portion of partially matching PacBio sequence plus the length of diverged full-length matches). In the Illumina data, the D estimates obtained from the frequencies in Figure 4 are much smaller, lying between 7% and 20% of ML.

This enrichment could be due to the greater retention of mitochondrial sequence (fraction M) in the preparation of the short-read libraries. Short-read libraries are usually fragmented and size selected to produce a distribution of fragments around 350-550 bp long, which will include fragments of the mitochondrial genome. On the other hand library preparation for long-read sequencing involves more careful shearing and a subsequent size selection for longer fragments (around 3-4 kbp in this case). This may cause mitochondrial genomes, starting at 16kbp before shearing, to be differentially lost from PacBio libraries while Numts, being part of the nuclear DNA, would be represented in their natural proportion.

### Estimating the genomic proportion of Numts

Building on the results presented above, there are two ways of estimating the genomic proportion of Numts in genome skimming data, which are possible even in the absence of a genome assembly. It is shown in Supplemental Information S6 that for our short-read data, there is a good correlation (R²=0.93) between the proportions of Numt-derived data reads (fraction D) and non-ML data (fraction A). The slope of this regression corresponds to the estimated genomic proportion. It is 0.09 %. This is a lower-bound estimate, because it is based on sequence divergence between Numts and the mitochondrial genomes sequence. The estimate based on PacBio CCSs is somewhat lower; ML CCSs with divergent sequences amount for 0.01% of the PacBio data. Another class of CCSs, which match only along part of their sequence, are likely to represent Numts, too, however there is a small chance that some of them are derived from chimeric SMRT bells. These sequences amount for 0.027% of the PacBio data, giving a total of 0.037%.

### Genome size and Numt content

*P. pedestris* has the largest genome of any insect listed in the Animal Genome Size Database (Gregory, 2016, accessed 24 June 2019). Its C-value of 16.93 corresponds to 16.5 Gbp (Doležel, Bartoš, Voglmayr, & Greilhuber, 2003). Consequently, a genomic proportion of just under 0.1 % is equivalent to about a thousand full length mitochondrial genomes inserted into the nuclear DNA (this length of sequence is equivalent to an entire *Drosophila melanogaster* or *Arabidopsis thaliana* chromosome). Although this is a surprisingly large number, as a proportion of the total genome it is consistent with estimates from other species. Hazkani-Covo et al. (2010) present estimates of Numt contents for a diverse list of 85 species ranging from 0 to 0.25% in multicellular organisms. By contrast RepeatExplorer analyses suggest repeats account for approximately 70 % of the *P. pedestris* genome, including transposable elements able to excise and re-insert themselves, providing a mechanism for copy-number increase.

### Tangle plots

In Fig. 5, we show tangle plots, which allow visual comparisons between Numts in multiple samples. The repeated occurrence of the same links within a sample suggest that rearranged mitochondrial sequence has been replicated within a single genome (the low coverage of < ⅓x makes repeated sequencing of the same region unlikely). The similar patterns in different individuals and populations suggest that many of these replicated insertions are fixed or occur at a high frequency. Given that they are unlikely to be functional, it is most plausible that they have spread by genetic drift.

Tangle plots can be used to visualise any short-read data sets mapped to a circular (or tandem-repetitive) reference, see Supplemental Information S7.

## Supporting information

Archive with supplementary information (GZ-compressed TAR ball)

## Acknowledgements

This research utilised Queen Mary’s MidPlus computational facilities, supported by QMUL Research-IT and funded by EPSRC grant EP/K000128/1. Further computational resources were provided by the ELIXIR-CZ project (LM2015047), part of the international ELIXIR infrastructure. HB was funded by a PhD studentship of QMUL’s School of Biological and Chemical Sciences. We wish to thank Graham Stone and Duncan Greig for their comments on an earlier version of the manuscript. We would also like to thank Alex Twyford for suggesting the name “tangle plot”.

## Data Accessibility

Mito sequences aligned – Supplemental data S1

Neighbour-joining tree – Supplemental data S2

Distances of paired reads mapped – Supplemental data S3

RE contigs – Supplemental data S4 matching PacBio reads, SAM format, GZIP-compressed TAR ball – Supplemental data S5 Regression plot – Supplemental data S6 Information on tangle plots – Supplemental data S7 (see also GitHub: https://github.com/SBCSnicholsLab/tangles)

The original sequencing data will be deposited on Dryad.

## Author Contributions

HB and RAN conceived the experiment and collected samples. HB carried out the experiment, analysed the data, and wrote the manuscript. HB and RAN revised the manuscript.

