## Supplementary material for "Mitochondrial sequences or Numts – By-catch differs between sequencing methods": Archive with supplementary information (GZ-compressed TAR ball): SI6_regression.docx

### S6 – Regression of the amount of non-mitochondrial-like data on the proportion of diverged Numts (D) in the whole data set

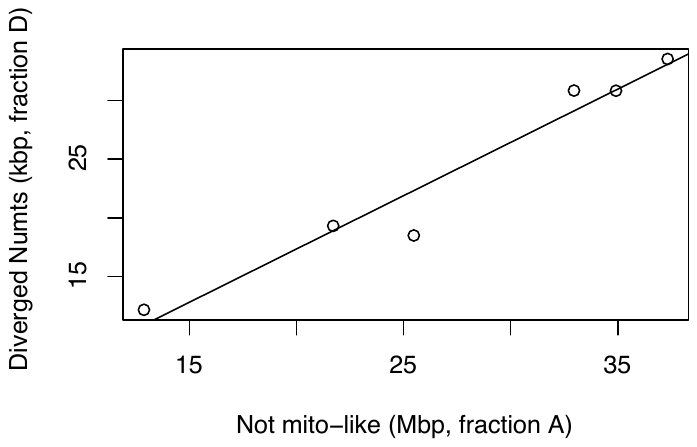

Sequence content of diverged Numts plotted against the fraction of not mito-like sequences. There is a good correlation (R^2^=0.93). The slope allows the estimation of the genomic content of Numts.

```
Coefficients:
              Estimate Std. Error t value Pr(>|t|)
(Intercept) -8.087e+02  3.069e+03  -0.264  0.80516
a            9.077e-04  1.065e-04   8.525  0.00104 **
---
Signif. codes:  0 ‘***’ 0.001 ‘**’ 0.01 ‘*’ 0.05 ‘.’ 0.1 ‘ ’ 1

Residual standard error: 2214 on 4 degrees of freedom
Multiple R-squared:  0.9478,	Adjusted R-squared:  0.9348
F-statistic: 72.67 on 1 and 4 DF,  p-value: 0.001039
```
