## Supplementary material for "Mitochondrial sequences or Numts – By-catch differs between sequencing methods": Archive with supplementary information (GZ-compressed TAR ball): SI7_Supplemental information X Tangle Plots.docx

Tangle plots are a way of visualising paired reads mapped to a circular reference. In the situation presented here (mapping genomic and mitochondrial sequences to a mitochondrial genome assembly), the reads of most pairs are expected to map in close proximity, because the sequencing libraries’ insert sizes are much shorter than the circular reference. This causes a pattern of short tangles following the circular outline of the plot. These are usually not plotted. More interestingly, there are some tangles, which go across the circle, evidence for nuclear insertions of mitochondrial origin.

### Implementation

Tangles are plotted with an R script, which takes as input a gz-compressed text file generated from a BAM file (sorted by name). The text file has to contain three tab-separated columns: Read name, mapping position, and BWA mapping quality like this example:

```
NS500784:18:H7MKYBGXX:1:11101:10138:1862	14558	AS:i:150
NS500784:18:H7MKYBGXX:1:11101:10138:1862	14913	AS:i:151
NS500784:18:H7MKYBGXX:1:11101:10757:10236	2715	AS:i:151
NS500784:18:H7MKYBGXX:1:11101:10757:10236	2969	AS:i:151
NS500784:18:H7MKYBGXX:1:11101:11422:18066	13054	AS:i:151
NS500784:18:H7MKYBGXX:1:11101:11422:18066	13365	AS:i:149
NS500784:18:H7MKYBGXX:1:11101:11491:16022	13754	AS:i:112
NS500784:18:H7MKYBGXX:1:11101:11491:16022	14078	AS:i:116
NS500784:18:H7MKYBGXX:1:11101:11871:8101	11262	AS:i:151
NS500784:18:H7MKYBGXX:1:11101:11871:8101	11669	AS:i:149
```

#### Software requirements

- BWA: for read mapping
- Samtools: for sorting and extraction of information from the BAM file
- R (v3.4.4 tested)

#### Instructions

1. Read mapping should be carried out in single-end mode because, when mapping paired reads, BWA tries to estimate the insert size from the data and will preferentially place reads at a “reasonable-looking” distance. The output should be a BAM file.
2. The BAM file needs to be sorted by read name. (samtools sort -n -@ <no of threads> -o <output.bam> <input.bam>)
3. Read names, positions, and mapping quality can be extracted with samtools and can be processed with GNU command line tools. (samtools view <infile> | cut -f 1,4,14 | gzip > <outfile.gz>)
4. The actual plots are generated interactively in R with the script tangles.R. The script contains code to re-plot tangle plots in the paper (figure 5).

### Other applications for tangle plots

Tangle plots are not restricted to showing mitochondrial data, any mapping of paired reads to a circular (or repetitive) sequence may be represented in a tangle plot

#### Podisma pedestris rDNA

The figure below shows a tangle plot generated from whole genomic DNA mapped against a rDNA reference (approx. 25 kbp).

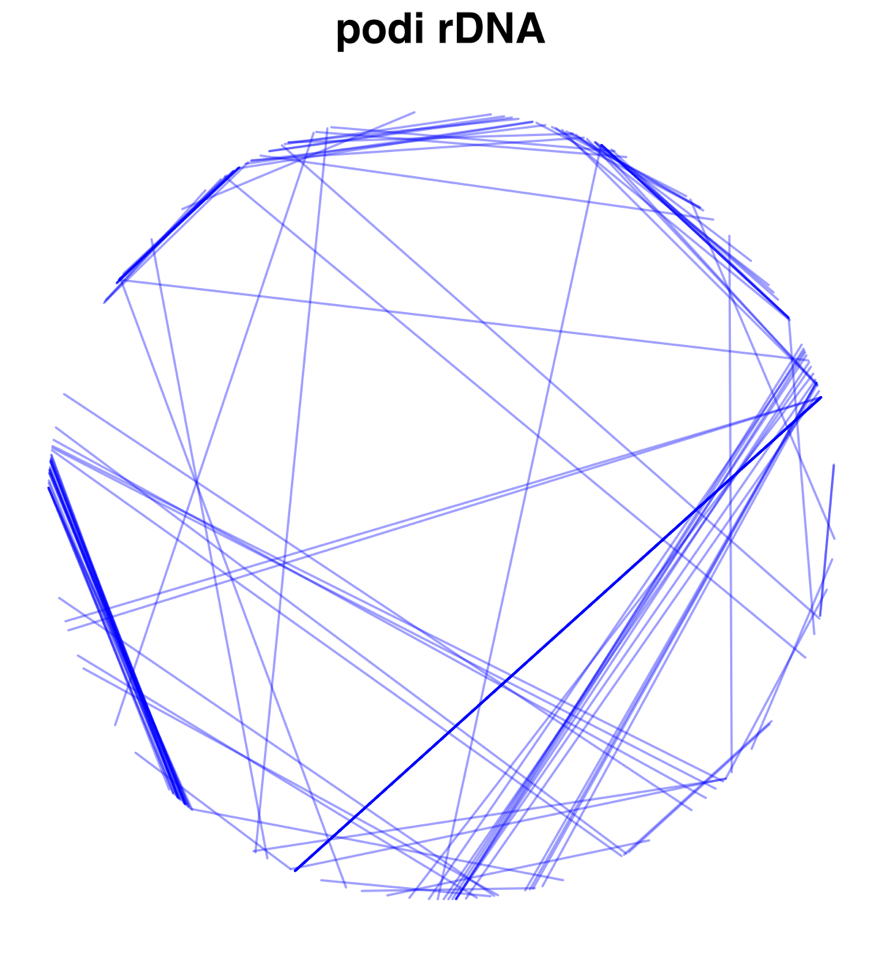
