## Supplementary figures and images for "Mitochondrial sequences or Numts – By-catch differs between sequencing methods"

### SI2_tree.pdf

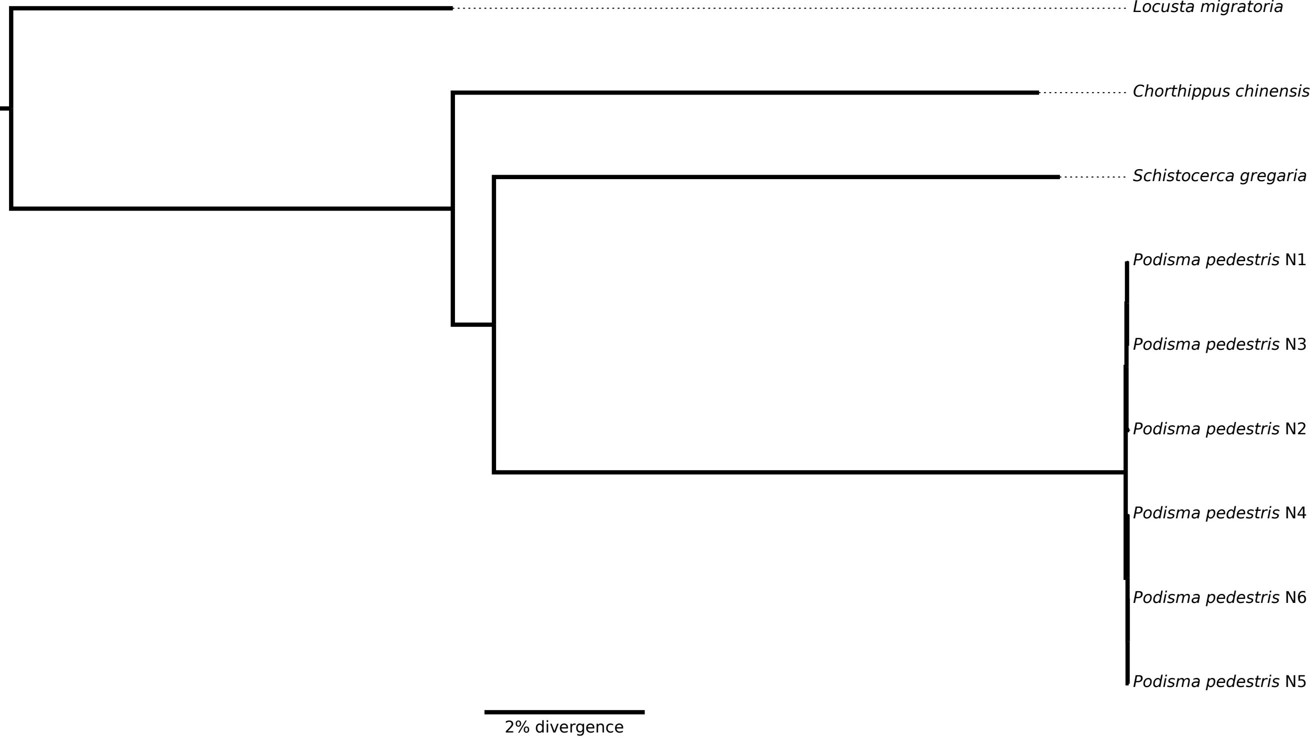

### SI3_insert_sizes.pdf

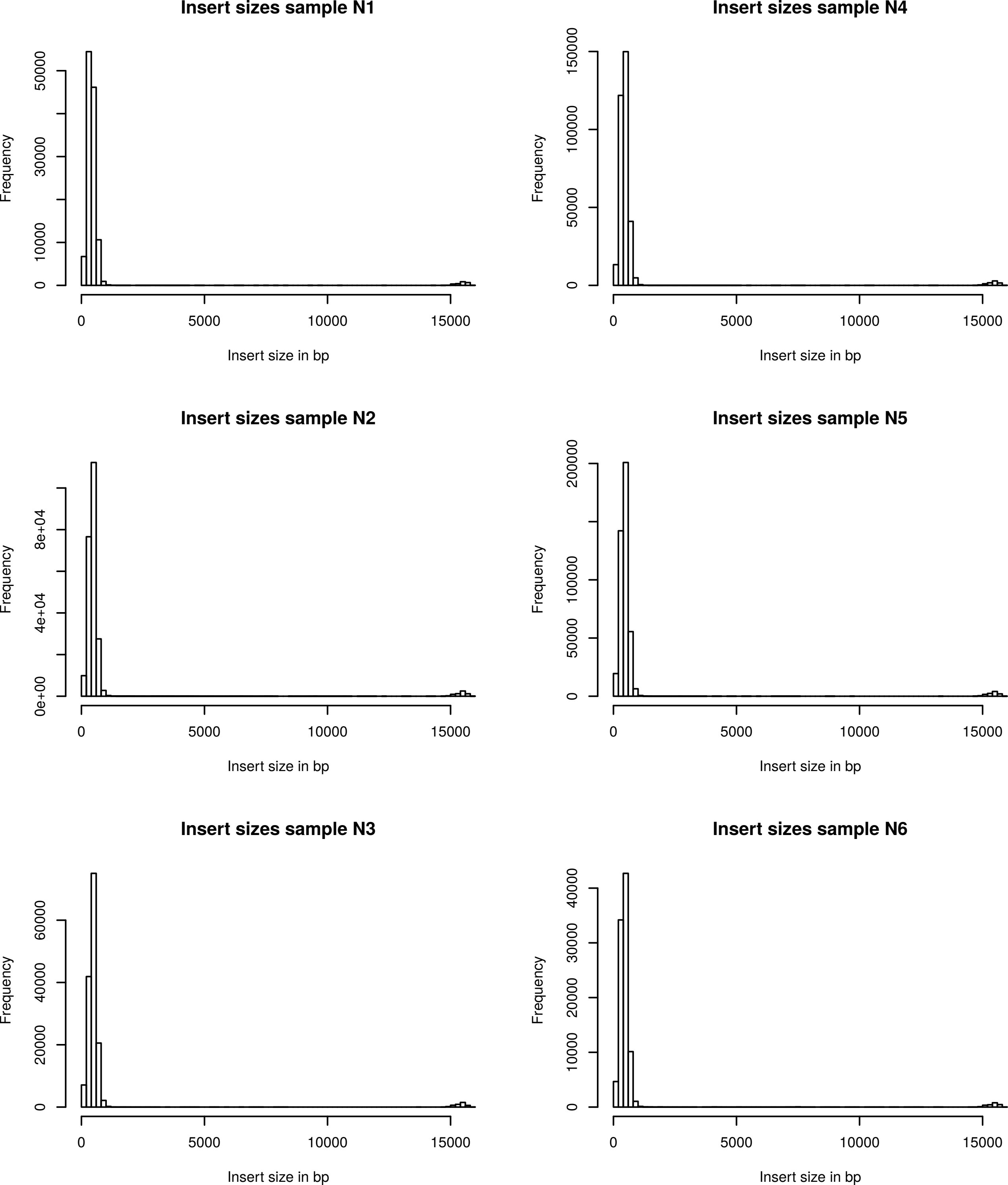
